# Expansive Linguistic Representations to Predict Interpretable Odor Mixture Discriminability

**DOI:** 10.1101/2022.04.11.487927

**Authors:** Amit Dhurandhar, Guillermo A. Cecchi, Pablo Meyer

## Abstract

Language is often thought as being poorly adapted to precisely describe or quantify smell and olfactory attributes. In this work, we show that semantic descriptors of odors can be implemented in a model to successfully predict odor mixture discriminability, an olfactory attribute. We achieved this by taking advantage of the structure-to-percept model we previously developed for monomolecular odorants, using chemical descriptors to predict pleasantness, intensity and 19 semantic descriptors such as ‘fish’, ‘cold’, ‘burnt’, ‘garlic’, ‘grass’ and ‘sweet’ for odor mixtures, followed by a metric learning to obtain odor mixture discriminability. Through this expansion of the representation of olfactory mixtures, our *Semantic* model outperforms state of the art methods by taking advantage of the intermediary semantic representations learned from human perception data to enhance and generalize the odor discriminability/similarity predictions. As 10 of the semantic descriptors were selected to predict discriminability/similarity, our approach meets the need of rapidly obtaining interpretable attributes of odor mixtures as illustrated by the difficulty of finding olfactory metamers. More fundamentally, it also shows that language can be used to establish a metric of discriminability in the everyday olfactory space.

## Introduction

Language is often thought as being poorly adapted to precisely describe or quantify smell and in particular olfactory properties such as the similarity or the discriminability of two molecular mixtures. Contrary to this idea, we have previously shown that for pure odors, it is possible to build models using the chemical structure of molecules [1] to predict the perceptual values of natural language attributes of smells [2]. Also, older studies have shown that a direct comparison of pure odor profiles based on 146 semantic descriptors [3] or down to 25 semantic descriptors [4], can be used to generate a distance between two pure odors that is highly correlated with their similarity ratings (*r* > 0.85). In these studies several metrics, such as Euclidean and Chi-squared distances or Tversky similarity, were used and the latter one was found to better match the similarity between two pure odors [4]. These past results show that indeed semantic descriptors can be used to quantify smell attributes, at least in the reduced universe of monomolecular odors.

Odor mixtures, the real-world situation for olfactory perception, add an extra level of complexity because the overall perceptual qualities of a mixture are not the sum of the qualities of the molecules composing it. This has led to, in our opinion, the erroneous suggestion that perception in olfaction, as for vision and audition, is object-oriented and that odor-objects are buried in such odor mixtures [5, 6]. Recent studies have shown that a model using a simple set of 21 chemoinformatic structural descriptors can predict with a significative correlation the odor similarity and discriminability between mixtures of molecules (*r* = 0.31 to *r* = 0.51 for both equi-intense odors and with varying intensities) [7]. In this work, we show that, besides chemoinformatic features, semantic descriptors of odors can also be implemented in a most effective model to predict odor mixture discriminability and similarity. This allows, through analysis of the semantic relationship between olfactory attributes, an interpretation of why 2 mixtures of molecules smell similarly, besides having components in common. We do this by taking advantage of the structure-to-percept model we previously developed for pleasantness, intensity and 19 semantic descriptors describing monomolecular odorants [1], to build a model able to predict the discriminability between any two odor mixtures as measured by a triangle test (see Figure 1). It not only meets the need of rapidly obtaining interpretable predictions of odor mixture discriminability, but also shows that language can be used to establish a metric in the everyday olfactory space.

**Figure 1:**
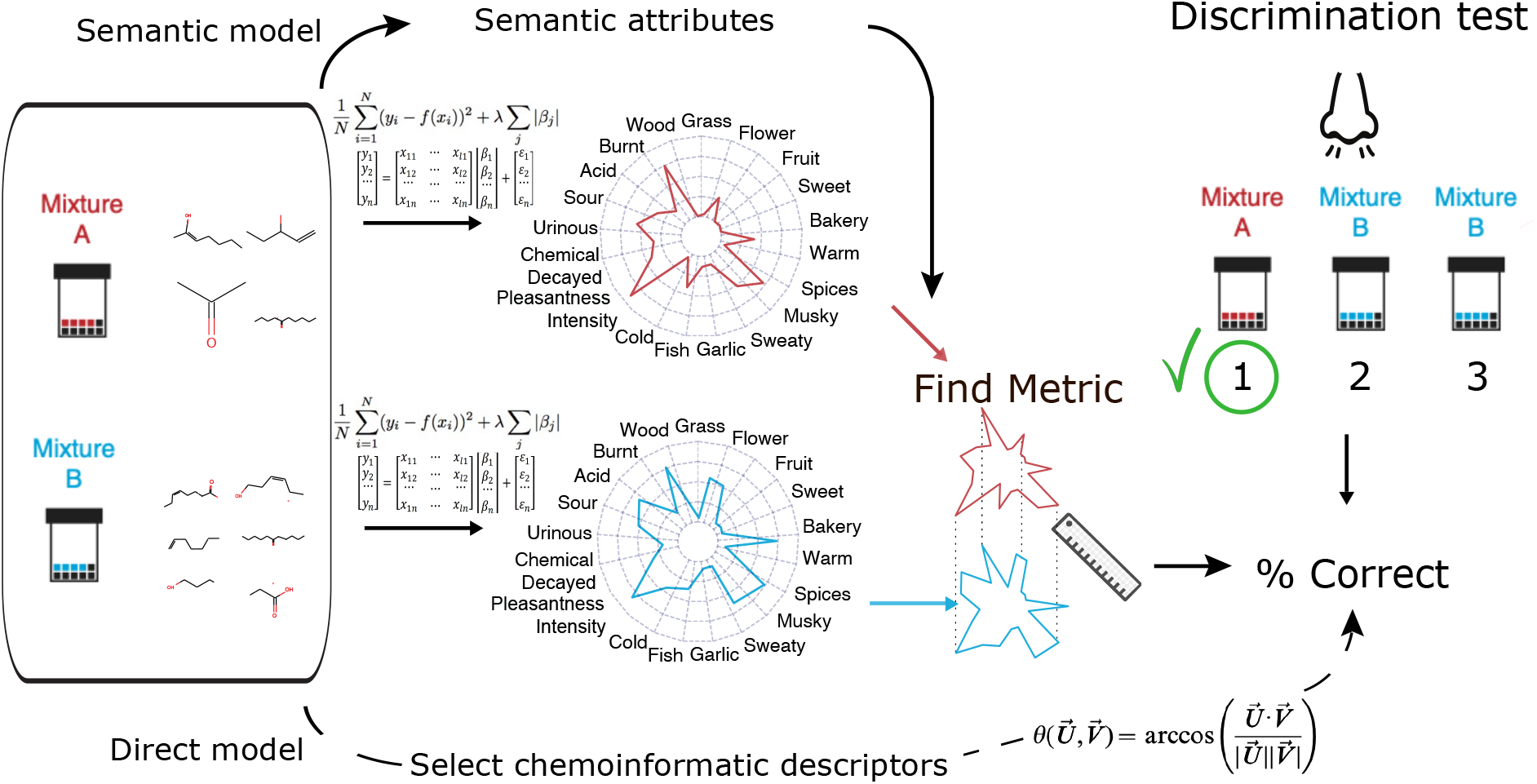
Predicting discriminability of odor mixtures. Diagram showing two possible solutions to predict the results of a discriminability test (*Right*) between two odor mixtures composed of different molecules (red and blue). A discrimination test consists of identifying one of the 3 vials that has a different odor and is quantified as the fraction of subjects that perform this task correctly. *Top*. Our solution consists on implementing a model that predicts semantic attributes of pure molecules from chemoinformatic features and then perform a metric learning to obtain the discriminability value. *Bottom*. The existing solution consists on directly predicting discriminability using for each mixture the values of 21 selected chemoinformatic features to calculate an angle distance between the two vectors. In both models the chemoinformatic features from different molecules composing the mixture are integrated linearly.

## Results

### Interpretable metric learning from semantic attributes

Instead of taking a direct approach to predict odor mixture discriminability from chemical descriptors [8, 7], we decided to implement a model with an intermediary step that expands the dimensions using semantic descriptors [1, 2] (see Figure 1). Discriminability experiments consist of finding the percentage of subjects that can discriminate the odor that is different when presented with 3 vials where 2 have an identical odor mixture, sometimes diluted [9, 6]. The expansion was done taking advantage of a model we previously developed [1, 2] to predict, from the chemoinformatic structural descriptors of any molecule, the values for intensity, pleasantness and the 19 semantic descriptors (see Figure 2a Top). We then averaged each of these 21 values across all the molecules in the mixture, irrespective of their number, to obtain the perceptual values of the mixture. The underlying approximation for this being that the perceptual contribution of each molecule to the olfactory mixture is independent. Given the clear limitations of this approximation and instead of trying several metrics as previously done [4], we decided to perform metric learning [10] to match the defined distance between the predicted perceptual values of any two mixtures to their experimentally measured discriminability. We implemented a *Mahalanobis distance*, that is a weighted *Euclidean distance* between the semantic descriptors of the two mixtures in the olfactory perceptual space. Namely, the *Mahalanobis distance* Δ between ***x**^i^* and ***y**^j^* is given by Δ^2^ = (***x**^i^* – ***y**^j^*)^⊤^Σ^-1^(***x**^i^* – ***y**^j^*), Σ is a covariance matrix that we chose to be diagonal to allow interpretability of the fitted weights. The initial unit value for each descriptor weight was changed and fitted in order to predict the fraction of subjects that can discriminate a given pair of odor mixtures obtained from an experimental dataset (see Figure 1 and Methods). Given that in Bushdid *et al* subjects discriminated up to 260 intensity-matched different odor mixtures composed of 10 to 30 molecules and varying the amount of shared components (see Methods), it seemed like the perfect dataset to perform metric learning (see Figure 2a Middle). The weights obtained for ‘intensity’, ‘pleasantness’ and the 19 semantic descriptors when performing a 10-fold cross validation scheme for the metric learning are shown by decreasing importance in Table 1. As we performed a Lasso regression to obtain the best metric fit using a minimal set of descriptors, 10 of the 21 weights are null and those descriptors do not contribute to the metric (see Table 1). The regression was also performed with a constant term, defining a lower limit for the distance, whose value was 0.5185. The good performance of this metric learning as measured by Root Mean Square Error (RMSE) can be seen in Figure 2b for Bushdid and 3 other different datasets.

**Figure 2:**
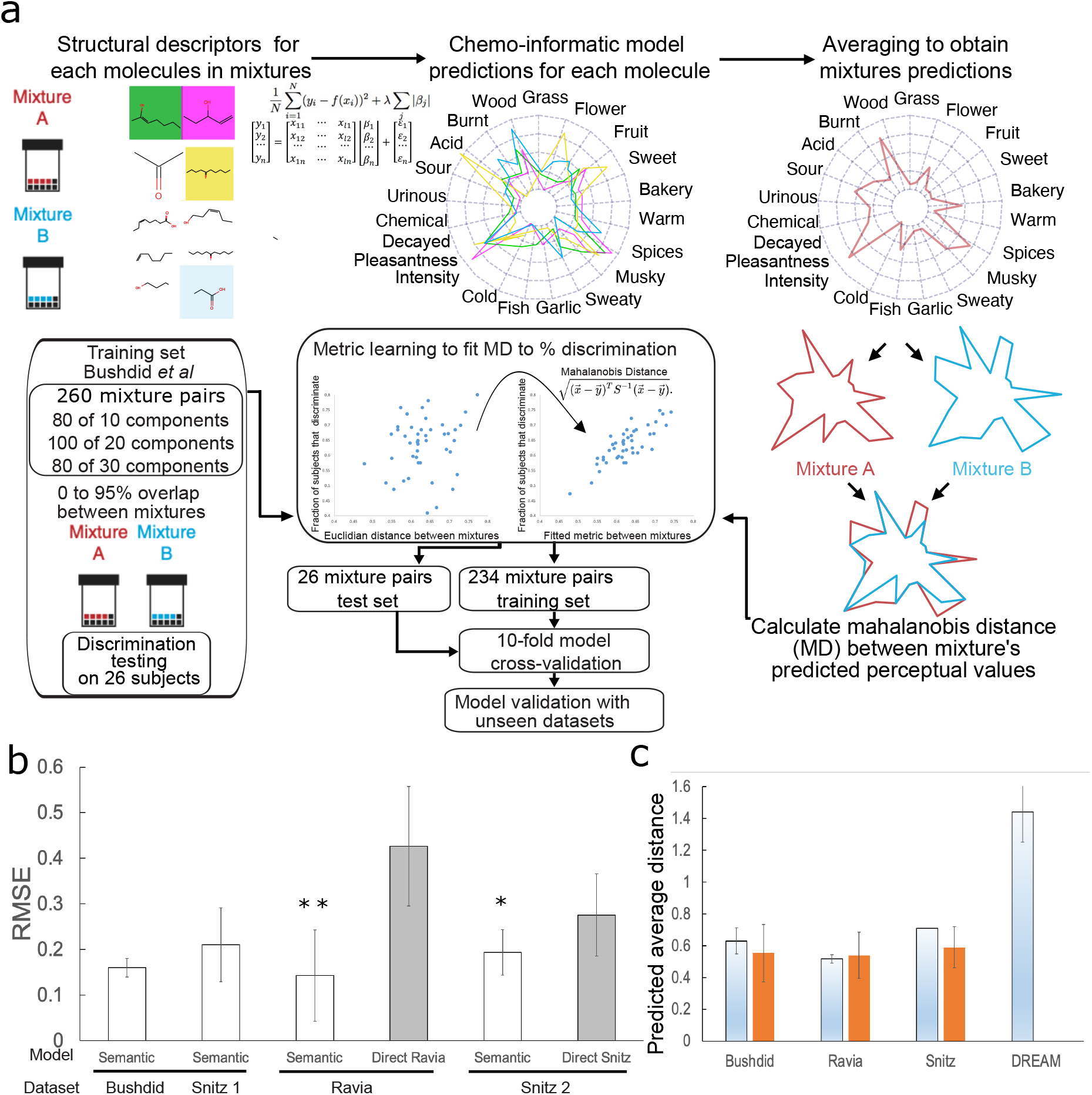
Metric learning of the Semantic Model for odor mixture discriminability. **a.** Diagram showing the flow of the metric learning for odor mixture discriminability. **b.** RMSE of the predictions of the *Semantic* model using semantic descriptors and the learned metric (white bars) for different datasets (shown below horizontal black bars) compared to the *Direct* model (grey bars). Significance indicated by *=p< 0.01 **=p< 0.001. **c.** Comparison of the predicted discriminability (light blue gradient bars) and the experimentally measured discriminability (orange bars) for 3 different datasets. Discriminability for DREAM dataset was not measured.

**Table 1:** Weight values for Mahalanobis Metric.

| Semantic descriptor | Weight Value | Semantic descriptor | Weight Value |
| --- | --- | --- | --- |
| FISH | 0.0497850544197713 | SPICES | 0 |
| COLD | 0.0347755400391788 | WARM | 0 |
| SOUR | 0.0169828569816300 | SWEATY | 0 |
| BAKERY | 0.0138228143227150 | WOOD | 0 |
| GRASS | 0.00848871765378457 | FRUIT/FLOWER | 0 |
| BURNT | 0.00430759902131422 | URINOUS | 0 |
| GARLIC | 0.00358620442521323 | MUSKY | 0 |
| SWEET | 0.00122436101055212 | ACID | 0 |
| INTENSITY | 0.000719940220699876 | PLEASANTNESS | 0 |
| CHEMICAL | 0.000107958448313051 | DECAYED | 0 |

In order to compare our approach to the previously published *Direct* model and validate it on unseen datasets, we implemented our *Semantic*-based model on 2 different datasets [8, 7]. The Snitz dataset consisted first of 147 pairs of intensity-matched different odor mixtures extracted from a total of 49 mixtures composed from 1 to 30 molecules (Snitz 1), and second of 91 pairs of intensity-matched different odor mixtures extracted from a total of 14 mixtures composed from 4 to 10 molecules (Snitz 2). In both cases we excluded experiments where the mixtures were the same. Although in these datasets the subjects were asked to rate the similarity between mixtures from 1 to 100, we obtained the discriminability as being 1 minus the similarity re-normalized between 0 and 1. The second external dataset was obtained from a discriminability experiment in a recently published article [7] where the odor mixtures did not have intensity matching and 50 pairs of different odor mixtures from a total of 120 mixtures composed from 10 to 30 molecules (Ravia). For all this heterogeneous external datasets our *semantic* model trained on data from Bushdid *et al* fared similarly and was much better than the *Direct* model whose RMSE was doubled (compare white and grey bars in Figure 2b.). Although we think that RMSE is the best metric when considering similarity or discriminability between odors, i.e their distance in the perceptual space, our results are comparable to the *Direct* model when using Pearson correlation as a metric (*r* = 0.333). The results of our *Semantic* model predictions are consistent when comparing the overall discriminability predictions to the actual experimental values for the Bushdid, Snitz and Ravia datasets (compare predictions grey bars and experimental results orange bars in Figure 2c.). Predictions also make sense when the model is used to obtain the discriminability between all possible pairs of the 476 monomolecular odors of the DREAM dataset [1], as given the diversity of molecules the average discriminability is expected to be much larger than for the other datasets (Figure 2c.).

### Using the semantic model to interpret properties of the olfactory space

Having a well performing validated model, we now interpret its implications. As observed in Table 1, the metric learning reduced the number of necessary attributes from 21 to 10 which are well distributed across the olfactory semantic space (see blue colored attributes on dendrogram in Fig.3). Not only that, but the 11 attributes that were discarded are either neighbors of another attribute in the dendrogram, i.e ‘grass’ and ‘wood’, or in the near vicinity, as if the Lasso regression was weeding out redundant terms as measured by their semantic similarity. This points to the relevance of using language to quantify the properties of smell. Also, it is interesting to note that 10 dimensions have come up in several studies as to seemingly underlie the structure of the olfactory space [11].

**Figure 3:**
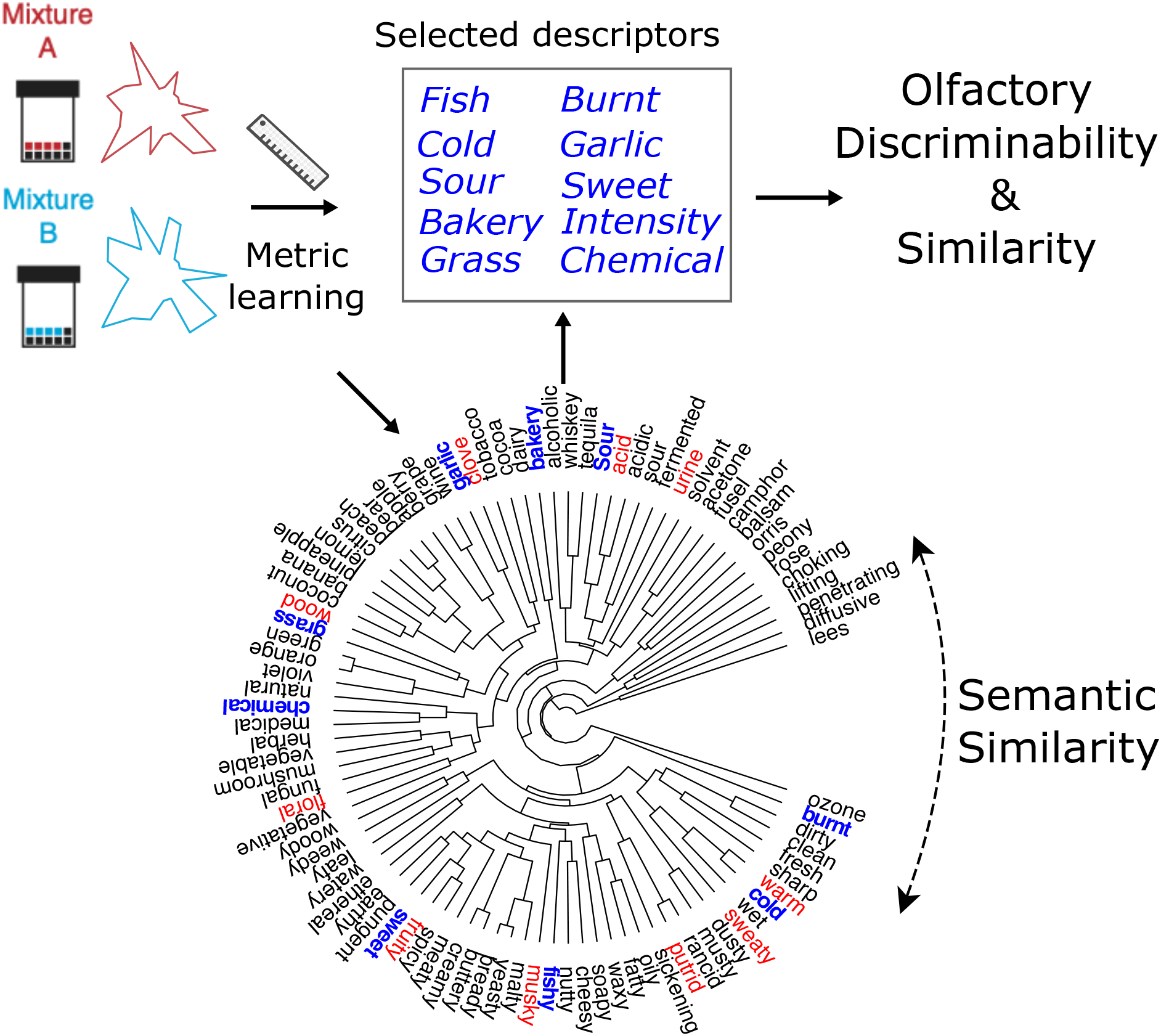
Language as a measure of smell. Diagram showing how the lasso metric learning for odor discriminability was able to extract ‘intensity’ and 9 optimal semantic descriptors out of 19, shown on table by order of importance. The dendrogram below shows 131 olfactory descriptors ordered by semantic similarity as measured by their cosine distance using vectors from word embbedings [12, 2] and in blue are whown the 10 optimal descriptors, equally distributed along the dendrogram. The 11 other red descriptors were eliminated during the optimization for being redundant, next to a blue descriptor, or not contributing to the metric. If the semantic attributes of mixtures used to describe them are different than the ones in blue, a transformation using the cosine distance between descriptors as used to construct the dendrogram can be implemented to project their values to the blue ones used in the model.

The semantic model can also be used to study the olfactory space in two dimensions: Exploring how the distance between 2 mixtures changes with the number of molecules and the overlap between the molecules composing them ((Fig.4a & b). Perceptual predictions from the 578 molecules (see Supplementary Data) that compose the Snitz, Ravia and DREAM datasets were used to calculate distances between: 1) a set of 100 randomly selected different pairs of mixtures for molecules with no overlap and where the number of molecules in the mixture were 1, 5, 10, 50 and 100 (inset Fig.4a) 2) a set of 100 pairs of mixtures composed of 10,20 and 50 randomly selected molecules with varying degree of overlap Fig.4a). Irrespective of the mixture composition, the average distance between mixtures decreases monotonously with the numbers of components towards what has been called an olfactory white [13], but also naturally, with the overlap between the mixtures. The decline is steeper for mixtures with less components and as the mixtures share less and less molecules (Fig.4a). Figure 4b synthesizes these results in a single 3 dimensional plot showing simultaneously how the distance between pairs of mixtures changes with the number of components and overlap between them. The region where mixtures can be distinguished is relatively reduced compared to the overall space of mixture possibilities.

**Figure 4:**
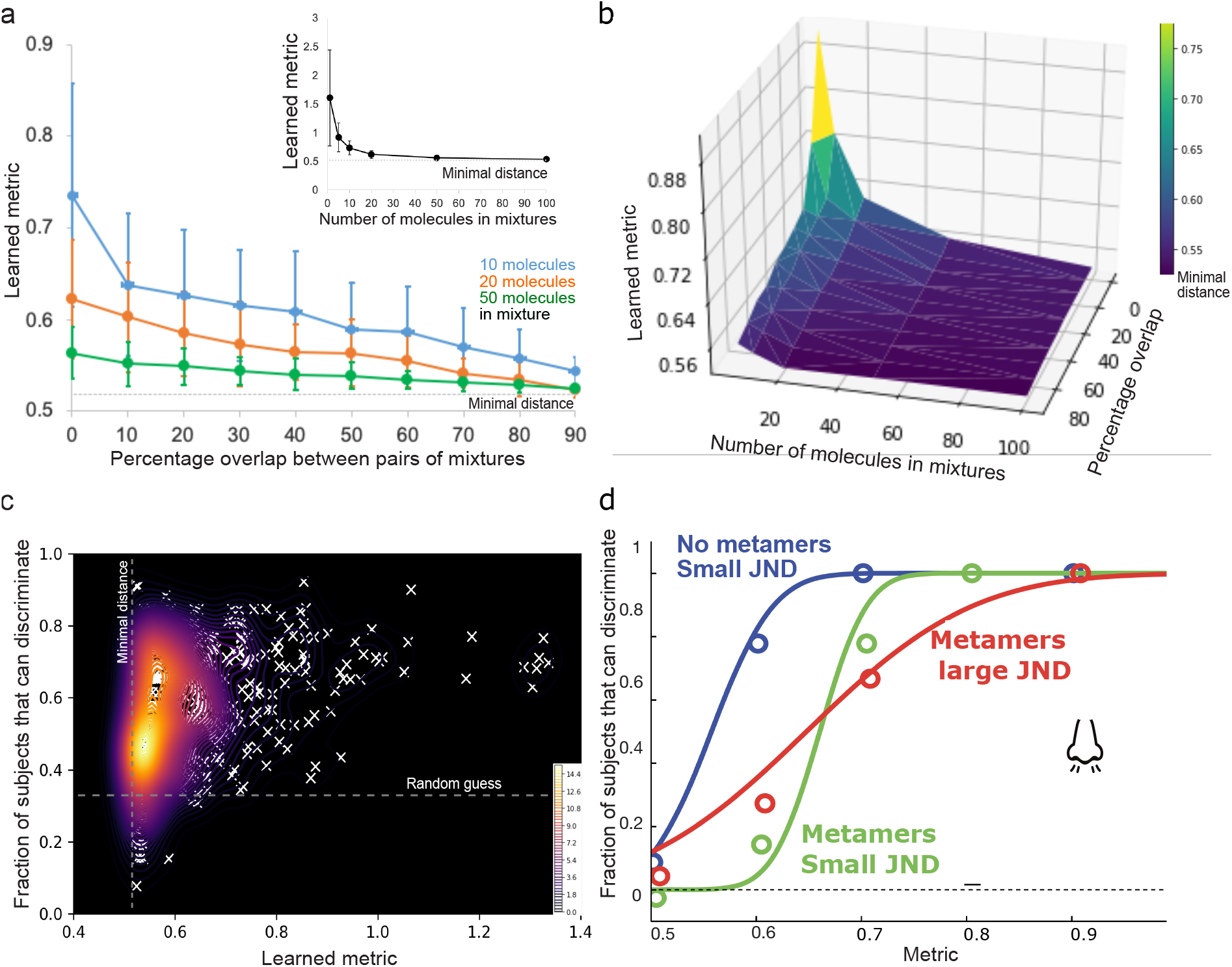
Metric-derived properties of the olfactory space. **a.** Plot showing the variation of the learned metric predictions between 100 pairs of odors for each point as the overlap between the components of the two mixtures varies. This was repeated for mixtures with 10 molecules (blue), 20 molecules(orange) or 50 molecules (green).Inset shows a similar plot of the variation of the learned metric predictions between 100 pairs of odors for each point as the number of elements in the mixtures varies and with no overlap between the components of the two mixtures. **b.** 3D plot of learned metric of mixture discriminability against mixtures overlap and number of components in a mixture. **c.** Density plot representing the 548 predictions (Learned metric) of the *Semantic* model for the 3 datasets against the experimental results for discriminability. Density colorbar is shown on lower right and single points are represented by white crosses. Plot is interpreted as a psychometric perceptual curve where most of the predictions lie on a very steep zone near the minimal distance between mixtures. Note that the model was trained to predict in the [0 1] interval, but its values for certain pairs of odors go above this limit. **d.** Diagram of the different possibilities of interpretation of the psychometric perceptual curves relative to the existence of olfactory metamers, i.e odors that smell the same but have no molecular components in common. Blue curve represents the interpretation of **c.** where no metamers can exist given that the lower tail in the perceptual curve is absent and the Just Noticeable Difference (JND) is small, green curve also has a small JND but a lower tail allows the existence of metamers and red is a curve with no lower tail but a large JND that allows the existence of metamers.

The abrupt transition from discriminable to indiscriminable mixtures in the olfactory space may be a clue to why it has been so difficult to prove/disprove the existence of olfactory metamers, i.e odors that smell the same but have no molecular components in common [7]. Hence, we next investigated in more detail the relationship between the *Semantic* model predictions for the learned metric and the actual discriminability values for the 3 datasets for a total of 548 predictions (see Supplementary Data). These are represented in Figure 4c as a density plot that can be interpreted as a psychometric perceptual curve where most of the predictions lie on a very steep zone near the minimal distance of 0.5185. The curve plateaus around a value of 0.8 and plunges around the minimal metric value, the zone of highest density, complicating the definition of a robust *Just Noticeable Difference* in olfactory quality, i.e the smallest distance between odors that are reliably discriminated (see also 4d). Indeed, the diagram of Figure 4d represents two different situations where metamers could exist and one were they do not. If the JND is small, as in our case, a lower tail in the perceptual curve is necessary in order to theoretically be able to find metamers (green and blue curves in Fig.4d). Indeed, if metamers exist in the transition part of the curve, dues to its steepness even a small difference between mixtures can generate a large perceptual change, making it difficult to find metamers. If the JND is large, then no lower tail is needed to allow the existence of metamers (red curve in Fig.4d) as they now can be found in the transition phase of the curve.

## Conclusion

We here describe a universal model to predict the discriminability and similarity between any pair of molecular mixtures. We achieved this by performing a linear integration of predictions of 21 olfactory attributes of pure odors using molecular descriptors to generate the percepts for odor mixtures and then fitting a Mahalanobis Distance to the discriminability score. This allowed to expand the description of the olfactory perception and only 10 of the olfactory attributes were needed to perform better than state of the art in several unseen datasets for the more relevant RMSE metric and comparably for Pearson correlation. We also show that as perceptual values can be converted, for any set of smell attributes, to the 19 semantic descriptors [2], the approach here described can be generalized. Overall, our results show that language can indeed be implemented in a general manner as a measure of smell attributes and can be used to explain properties of the olfactory perceptual space. For example our model shows that the difficulty to find olfactory metamers is due to the steepness, i.e small *JND*, and absence of tail of the olfactory perceptual curve (Fig.4a). This results prove that language can be used to establish an interpretable metric of odor discriminability in the everyday olfactory space, and hence opens the possibility to establish it as a more general tool to quantify the effect of known changes in olfactory perception in diseases such as Parkinson’s [14], Schizophrenia [15] and more recently COVID [16].

## Methods

The predictions and validations for the Interpretable Semantic Olfactory Mixture Discriminability model were generated with the following steps: 1) Determine structural properties for each molecular mixture using Dragon descriptors 2) Predict the ‘intensity’, ‘pleasantness’ and 19 semantic descriptors of each mixture 3) Perform the metric learning where Lasso regression was used with a Mahalanobis distance metric fit with a constant term. The Mahalanobis distance is learned on the correct discriminability fraction on the training dataset [9], so that the values are fit to the [0,1] interval. The Bushdid data consisted of mixtures of 10, 20, or 30 components drawn from a collection of 128 odorous molecules previously intensity-matched. In doubleblind experiments, 26 subjects were presented with three odor vials, two of which contained the same mixture, whereas the third contained a different mixture. Each subject completed the same 264 discrimination tests. These tests result in a percentage of individuals (the right fraction) correctly distinguishing two mixtures. 4) Predict using 10-fold cross-validation on the Bushdid dataset [9] and on two previously published datasets ([8] and [7]). The Snitz dataset [8]consisted of 147 pairs of intensity-matched different odor mixtures extracted from a total of 49 mixtures composed from 1 molecule to 30 molecules (Snitz 1), the second Snitz dataset consisted of 91 pairs of intensity-matched different odor mixtures extracted from a total of 14 mixtures composed from 4 to 10 molecules (Snitz 2). Experiments where the mixtures were the same were exluded. Although in these datasets the subjects were asked to rate the similarity between mixtures from 1 to 100, we obtained the discriminability as being 1 minus the similarity re-normalized between 0 and 1. The Ravia dataset was obtained from a discriminability experiment in a recently published article [7] where the odor mixtures did not have intensity matching and 50 pairs of different odor mixtures from a total of 120 mixtures composed from 10 molecules to 30 molecules (Ravia). Note that the model was trained to predict in the [0 1] interval, but its values for certain pairs of odors go above this limit.

We now describe the details of implementation of the *Semantic* model. Predictions for the *Direct* model where kindly shared by Aron Ravia.

### Interpretable Olfactory Mixture Discriminability Method

In our method we want to learn an interpretable yet accurate metric that can map the similarity/discriminability between two odor mixtures in the semantic space to the experimentally measured human discriminability. Thus our method consists of the following steps:

1. Obtain semantic descriptors for odor mixtures. This could be done by using the olfactory models proposed in [1] to obtain mono-molecular predictions. The mono-molecular predictions of the molecules composing a mixture are averaged following each of the 21 dimensions to obtain the perceptual prediction for the mixture.
2. Compute model features defined as the squared difference between the corresponding perceptual values of the semantic descriptors of two mixtures. Formally, if [*x*_1_,…, *x_k_*] and [*x*_1_,…, *x_k_*] are the *k* semantic descriptors of the two mixtures respectively, then the *i*^th^ new feature *f_i_* = (*x_i_* – *y_i_*)^2^.
3. Train a lasso model 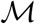 [17] that optimally maps the [*f*_1_,…, *f_k_*] to a target *D* that measures the human discriminability for a set of pairs of olfactory mixtures. The training is done on a dataset that has multiple mixtures with difficulty in distinguishing pairs of mixtures quantified for some subset.
4. Output the model 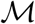.

Note that the above model is interpretable since the features *f_i_* correspond to specific the semantic descriptors that define *x_i_* or *y_i_* corresponding to the same descriptor *i.e* “fruity”. Moreover, the trained model is a sparse linear model and so weights are assigned to (few) individual *f_i_*s making the final output easy to interpret. In a certain sense, we learn a shifted Mahalanobis distance, as the bias term may be non-zero, quantifying the human discriminability between the mixtures. The design choices make the process interpretable and as shown in (Fig.4c), the semantic descriptors are selected in order to cover the semantic space and avoid redundance: Between “warm” and “cold” only the latter is chosen, similarly between “grass” and “wood”. A more general model would not necessarily maintain this interpretability such as when using a complex non-linear model or neural network instead of lasso, This would also happen if the new features were constructed as a concatenation of the original semantic features leading to a large dimensional feature space, then used for model training.

## Acknowledgements

We thank Aaron Ravia for sharing his predictions and other materials, Pablo Polosecki for extensive discussions, the editor and reviewers for their useful comments.

